# Integration of a smooth mesh-based contact pressure model into tracking and predictive simulations

**DOI:** 10.64898/2026.09.14.751442

**Authors:** Mohanad Harba, Gil Serrancolí

## Abstract

Musculoskeletal simulations are widely used to estimate joint loading, yet most musculoskeletal models estimate knee contact forces as resultant forces or as normal medial and lateral contact forces, without resolving pressure distributions across the articular surfaces. This paper presents a smoothed mesh-based knee contact pressure model that computes continuously differentiable tibiofemoral contact pressures, enabling its direct integration into full-body movement simulations. Built on an elastic foundation formulation, the model introduces smooth approximations ensuring that all contact functions and their derivatives remain continuous throughout the simulation. A systematic sensitivity analysis was performed across five key parameters: mesh resolution, joint damping and three smoothing parameters. Tracking simulations across eight gait trials demonstrated that the nominal configuration achieved mean RMSE values for medial and lateral knee contact forces of 51.6 N and 75.2 N, respectively, with a mean RMSE for joint angles of 1.45*^◦^* and (*r* = 0.97), converging in less than three hours on a standard computer. Mesh resolution was identified as the dominant factor that affected both accuracy and convergence, while damping variations had negligible influence. As a proof of concept, the model was also incorporated into predictive simulations, demonstrating that increasing the weight on the contact pressure term in the cost function leads to reduced tibiofemoral loading, particularly in the lateral compartment. The proposed formulation provides a computationally efficient and numerically robust framework for simulating knee contact mechanics within full-body musculoskeletal models.

**Author summary:** Knee osteoarthritis is a leading cause of disability worldwide, yet the internal forces acting across the joint during movement are difficult or impossible to measure directly in most individuals. Musculoskeletal simulations offer a powerful alternative, but existing models either oversimplify joint contact into a single resultant force or are too computationally demanding to embed in full-body movement optimizations. We developed a new knee contact model that computes how pressure is distributed across the joint surface in a mathematically smooth and computationally efficient way. This smoothness is crucial, since it enables the model to be used in musculoskeletal simulations with gradient-based solvers and automatic differentiation tools to solve optimal control problems. We validated the model across eight gait trials from a subject with a knee prosthesis, demonstrating accurate reproduction of measured knee contact forces and joint kinematics. We also demonstrated the model’s use in predictive simulations, where penalizing joint pressure in the cost function produced gait adaptations that reduced tibiofemoral loading, particularly in the lateral compartment. This work opens the door to computational tools that can evaluate surgical interventions, assistive devices, and rehabilitation strategies directly in terms of their effect on joint pressure distributions.

## Introduction

Musculoskeletal simulations have become essential tools for studying human movement and supporting clinical and biomechanical decision making. They estimate internal variables such as muscle forces, joint reaction forces, and tissue loading which are difficult to measure experimentally, but strongly influence how musculoskeletal disorders develop and how rehabilitation and treatment strategies should be designed [1]. Instrumented knee implants have enabled direct measurements of joint contact forces in subjects [2, 3], but these datasets are limited and cannot describe detailed joint mechanics such as how pressure varies across the joint surfaces or how forces are distributed [4–6]. Consequently, accurate computational models are essential to understand tibio-femoral loading, improve implant design, and support subject-specific clinical decision-making [7]. Most musculoskeletal models estimate knee contact forces as resultant forces or as normal medial and lateral contact forces [6, 8, 9]. These approaches are popular because they are computationally efficient and integrate well with full body simulations, but they do not provide information about how pressure is distributed across the articular surfaces.

More detailed methods, such as finite element methods, have been used to estimate local pressure and stress distributions [10]; however, incorporating these models into musculoskeletal simulations based on dynamic optimizations would be highly computationally demanding. There have been studies using elastic-foundation models integrated into mesh-based contact models, which speed up the computation of joint pressures [11, 12], mainly used for inverse problems. However, existing collision-detection algorithms are highly non-continuously differentiable, which hinders convergence in musculoskeletal simulations based on optimal control problems (OCPs). In addition, pressure estimations obtained from these models depend strongly on defining parameters such as mesh resolution, stiffness, and damping parameters. Prior work has shown that contact pressures, penetration depths, and computation times can vary substantially depending on how these parameters are selected [13, 14]. The novelty of the proposed contact pressure model lies in its integration into full-body simulations, including compliant foot-ground contact models [15, 16] and compliant muscle-tendon models, as well as in the ability to incorporate joint pressure values either in the cost function or as constraints. Furthermore, by formulating the problem as an OCP solved with a direct collocation method, state continuity is enforced across the entire gait cycle, which is not necessarily guaranteed in tracking simulations that solve the problem using static optimization. In turn, musculoskeletal simulations based on the resolution of an OCP (without joint contact pressure models) using gradient-based methods have been shown to be solvable in around 30 min on a standard computer [17, 18]. Those could be formulated as tracking or predictive simulations.

The purpose of this study is to develop and evaluate a mesh-based knee contact model capable of computing smooth pressure distributions across the tibial contact surface during musculoskeletal simulations. Building on the elastic foundation formulation of Bei and Fregly [11], the proposed model introduces continuously differentiable contact functions to compute penetration and pressure values. This enables its use with automatic differentiation tools in gradient-based OCPs, where the contact pressures and forces must be smooth and continuously differentiable functions of the model coordinates and velocities to ensure good convergence and numerical efficiency [19]. The present model estimates local pressures over the tibial surface, enabling a more detailed assessment of how loads are transferred across the joint. In addition, we perform a systematic sensitivity analysis to quantify how model parameters such as mesh resolution, smoothing parameters, and damping coefficients affect computational cost, and kinematics and knee contact force tracking accuracy. Overall, this work provides a contact modeling approach that addresses current limitations and supports the development of more accurate and computationally efficient musculoskeletal simulations of the knee.

## Materials and methods

### Experimental data

The experimental data used in this study came from the Knee Grand Challenge competition to predict in vivo knee loads [2]. The data include three-dimensional marker trajectories, knee contact forces and ground reaction forces during gait cycles, for an 88-year-old participant (mass: 65 kg, height: 166 cm) wearing a knee prosthesis on the right leg, which was reconstructed using subject-specific data derived from a computed tomography (CT) scan [2]. For the tracking simulations, eight gait trials (from left heel-strike to left heel-strike) were analyzed, consisting of two repetitions for each of the following movement types: normal walking, bouncy gait, forefoot strike gait, and treadmill walking (at 1 m/s and 1.4 m/s).

### Musculoskeletal model

A full-body OpenSim musculoskeletal model with 34 degrees of freedom (DoFs) was used, including all six DoFs at the right knee with the proposed contact model, and with only the flexion-extension knee DoF at the left leg. The lower limbs were actuated by 94 muscles spanning 20 DoFs, while the 8 DoFs of the upper limbs were torque-driven. The full-body model was used first to parameterize muscle tendon-length and moment arms [20]. Then, the model was applied to tracking simulations to evaluate the performance of the proposed contact modeling approach, and to a predictive simulation to assess its effect on the predicted motion.

### Knee contact pressure model

A smoothed mesh-based contact model was developed to estimate contact pressures at the interface between the femoral and tibial components of a prosthesis. The main novelty is that the collision detection algorithm is smooth (continuously differentiable), which facilitates its use for gradient-based optimization using automatic differentiation. The geometry of both components is represented by triangular surface meshes. First, the contact geometries of both components (femoral and tibial) were simplified while preserving the relevant geometric detail (Fig 1A). To analyze the impact of mesh resolution on an OCP, we considered three discretizations: 171 femoral faces *×* 49 tibial faces (*∼* 54 mm^2^ per face), 258*×*75 (*∼* 28 mm^2^ per face) and 342*×*100 (*∼* 22 mm^2^ per face). Before running the simulations, we generated the set of candidate contact pairs using the experimental kinematics of a walking trial (see example in Fig 1B). At each time frame, each tibial face was paired with those femoral faces whose centroids lay within a 1 cm radius sphere centered on the tibial-face centroid. For this step, the knee was modeled as a pin joint.

**Fig 1.**
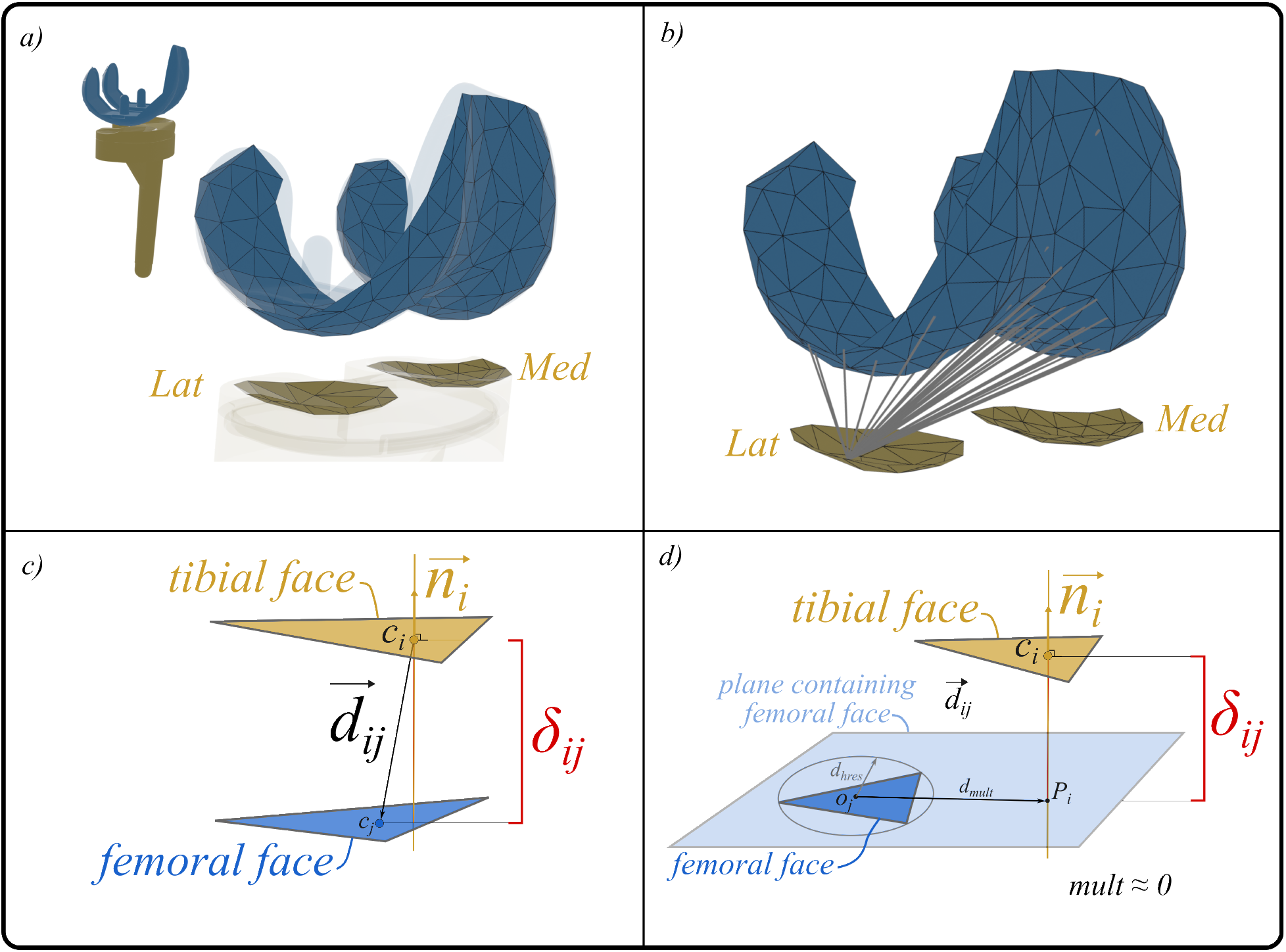
Prosthesis model and penetration calculation. A) Original prosthesis CAD model shown transparently and one of its simplified representations (75 faces for the tibial component and 258 faces for the femoral component). B) Example of potential contact pairs of one tibial face associated with all candidate femoral contact faces. C) Illustration of the penetration *δ_ij_* computation, where 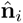 is the unit normal vector of tibial face *i* and 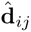 is the vector connecting the face centers. D) Example of a false-positive contact (*mult ≈* 0).

The contact pressure computation consists of the following steps:

**Step 1.** At each optimization iteration, the model computes the maximum normal penetration for each tibial face relative to its potential femoral face in contact. For tibial face *i* with centroid **c***_i_* and unit normal 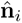, and for each candidate femoral face *j* with centroid **c***_j_*, we consider the displacement vector 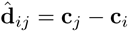 and project it onto 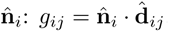. The penetration is defined as *δ_ij_*= *− g_ij_* (i.e., positive when penetration occurs along − 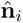) (see an illustrative example in Fig 1C).

**Step 2.** The maximum penetration *δ*_max_ for each tibial face is calculated using a smooth maximum (Mellowmax) function as defined in Eq (1, [20]) (see A.1), which converts the model continuously differentiable (note that 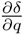 are continuous):

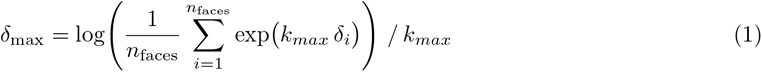

where *n*_faces_ is the number of candidate contact faces, *δ_i_*is the penetration depth of face *i*, and *k_max_* is the smoothing parameter.

**Step 3.** An elastic foundation model, described in [11], is applied to compute the contact pressure from the penetration value:

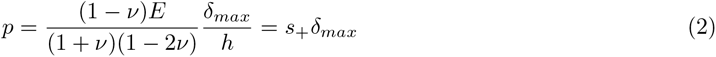

where *p* is the nominal contact pressure, *E* = 4 *·* 10^8^ Pa is Young’s modulus, *ν* = 0.46 is Poisson’s ratio, *δ_max_* is the maximum penetration depth as defined in Eq (1), and *s*_+_ is the effective stiffness relating *p* to *δ_max_* in the presence of penetration.

To provide the optimizer with gradient information in the vecinity of the contact transition, a small non-zero slope is retained when there is no penetration. Thus, we consider:

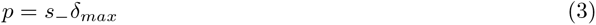

with *s_−_* = 10^6^ *Pa/m*. To limit the magnitude of non-physical negative pressures while maintaining a smooth transition between the two regimes, the final pressure expression is defined as:

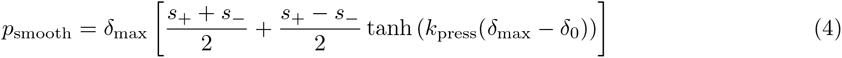

where *k*_press_ is a pressure smoothing parameter controlling the sharpness of the transition (see A.2 for details), and *δ*_0_ defines the location of the transition.

**Step 4.** To avoid false positive contact cases, a multiplier *mult* is introduced (see A.3). When a false-positive contact is identified, *mult* smoothly approaches zero; otherwise, it smoothly approaches one (see an illustrative example in Fig 1D). The details of the smooth formulation are provided in A.3.

**Step 5.** The final pressure is obtained as follows:

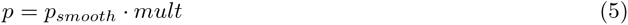

**Step 6.** Finally, the contact normal force vector for each face is computed as 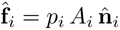, where *A_i_*is the area of the *i*-th face of the tibial component. The resultant contact wrench (a three-component force and a three-component moment) is computed at the center of the tibial component, and the corresponding reaction contact wrench at the femoral component. This information is transferred to compute the skeletal dynamics using the OpenSim API [21].

### Sensitivity analysis

To evaluate the sensitivity of the contact model to key parameters, we compared simulation results perturbing five parameters: the femoral-tibial mesh resolution (*n*_faces_); the smoothing coefficients for the maximum-penetration and pressure functions (*k*_max_ and *k*_press_); the multiplier transition parameter (*k*_ov_, which controls the sharpness of the transition between valid and invalid contact pairs, see A.3); and the knee-joint damping coefficients (*c*_F_, *c*_M_). Passive torques (*τ_pass_*) and forces (*F_pass_*) for the knee’s secondary degrees of freedom were modeled using a combination of non-linear stiffness and linear viscous damping terms:

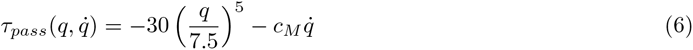

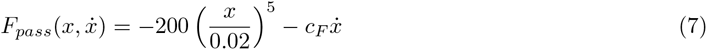

where *q* and *x* denote rotational (in degrees) and translational (m) coordinates, and 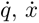 their corresponding velocities. These expressions are based on the assumption that the passive load increases when the coordinate moves outside its expected range of motion, similar to [17], with the damping coefficients *c_M_* and *c_F_* adjusted in the present work to examine the influence of viscous coefficients resistance on joint behavior and optimization convergence.

For each parameter, two alternative values were tested around a reference setting (hereafter referred to as ‘nominal’) defined by: *n_fem_* = 258 and *n_tib_* = 75, *k_press_* = 10000, *k_max_* = 10000, *k_ov_* = 100, *c_M_* = 0.001, *c_F_* = 1000. The two alternative values tested for each parameter are summarized in Table 1.

**Table 1.** Parameter values tested in the sensitivity analysis.

| Parameter | Case 1 | Nominal | Case 2 |
| --- | --- | --- | --- |
| $n_{\text{fem}}, n_{\text{tib}}$ | 171, 49 | 258, 75 | 342, 100 |
| $k_{\text{max}}$ | $2 \cdot 10^4$ | $10^4$ | *Max* |
| $k_{\text{press}}$ | $10^3$ | $10^4$ | $10^5$ |
| $k_{\text{ov}}$ | $10^3$ | $10^2$ | $10^4$ |
| $c_M, c_F$ | 0, 0 | 0.001, 1000 | 0.0005, 500 |

The nominal configuration is in the middle column. *Max* is the non-continuously differentiable maximum function.

These values were chosen based on a practical analysis of each parameter. The nominal mesh resolution (258*×*75) offered a suitable trade-off between geometric accuracy and computational cost. Finer meshes improved the spatial representation of contact pressures; however, they substantially increased runtime and memory use. The smoothing coefficient *k_max_* controls how closely the model approximates the true maximum penetration. Larger *k_max_* values provide a more accurate approximation but make the derivative 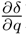 numerically stiffer. Conversely, smaller values produce smoother derivatives but yield a less accurate estimate of the true penetration. The parameter *k_press_* controls the smoothness of the pressure values: higher values make the transition from non-contact to contact steeper, whereas lower values yield a softer, more gradual contact pressure-penetration response. The parameter *k_ov_* controls how sharply the multiplier *mult* moves from 0 to 1. Smooth values produce a gradual transition between active and inactive pairs, whereas larger values enforce a sharper cutoff.

All combinations were implemented in identical tracking simulations across the eight analyzed gait trials. For each simulation, we extracted four performance indicators: (1) computation time, (2) number of optimization iterations, (3) accuracy of knee contact forces (KCFs), quantified separately for the medial and lateral compartments using the root mean square error (RMSE) and Pearson’s correlation coefficient (*r*) between the simulated and experimental KCFs, and (4) joint kinematic accuracy, computed as the mean RMSE and *r* across all tracked joint angles. For each parameter, the results of the eight gait trials were combined and summarized using the median and the median absolute deviation (MAD) to describe the typical performance and its variability.

Instead of focusing on a single metric (such as the lowest KCF error), we assessed the overall balance between accuracy and computational efficiency. A configuration that produced slightly improved knee contact forces but required substantially longer computation time, or resulted in less accurate joint kinematics, was considered worse overall. Finally, each parameter variation was evaluated relative to the nominal reference case to determine whether it led to consistent improvements across all movements without introducing substantial drawbacks in other aspects.

### Tracking simulations

The tracking simulations were formulated as an OCP using direct collocation, following the formulation described in [17, 20]. The integral cost function simultaneously minimized the squared difference between simulated and experimental data and muscle effort. The tracked quantities included full body joint kinematics, ground reaction forces and moments, and medial and lateral knee contact forces. Muscle activations were minimized to resolve the muscle redundancy problem, and regularization terms were included to ensure smooth state and control trajectories. The proposed contact pressure model was integrated into the simulation, and the resulting medial and lateral contact forces from the mesh-based model were used as knee contact force outputs for comparison with experimental measurements. Each simulation covered one full gait cycle, from left heel-strike to left heel-strike, across eight trials representing the four mentioned movement types. A total of 88 tracking simulations were performed to carry out the sensitivity analysis described in the previous section.

### Predictive simulations

Predictive simulations were also formulated as OCPs using the same direct collocation framework, but without tracking experimental data. Instead, gait was generated by minimizing a cost function that combined metabolic energy expenditure, muscle effort, and a pressure-based term aimed at minimizing the maximum pressure value at the tibial compartments. Joint passive forces and moments, arm excitations, and the coordinates for the secondary degrees of freedom at the knee were minimized to reduce redundancy. The main difference with respect to published predictive simulations [17, 22] is the use of the pressure contact model and without the symmetry assumption. For regularization purposes, control accelerations, time derivative of muscle activations and normalized tendon forces were also minimized with a lower weight.

Periodicity constraints were imposed on all state variables, and an average forward speed was prescribed to 1.33 m/s. The nominal contact model parameter values used in the tracking simulations from the previous sensitivity analysis were adopted for these predictive simulations. The results of four predictive simulations with varying weights for the pressure penalization term are presented (*w* = 0, no penalization; *w* = 0.03; *w* = 0.3; and *w* = 3). The formulation of the simulations to reproduce the results presented here can be found at https://github.com/gilserrancoli/MeshContactModel_forMSKSim.

## Results

### Tracking simulations

All nominal tracking simulations converged successfully for all eight gait trials. The simulations required less than 3 hours of computation time and fewer than 1600 optimization iterations (see Fig 2 and Fig 3). They achieved mean RMSE for medial and lateral KCFs of 51.6 *±* 7.9 N and 75.2 *±* 4.1 N, respectively, as well as a mean joint-angle RMSE of 1.45*^◦^*, with a mean correlation coefficient of *r* = 0.97 (see Figs 2 and 3, and Table 2). These results show that the nominal configuration provides accurate KCF estimates and kinematic tracking.

**Fig 2.**
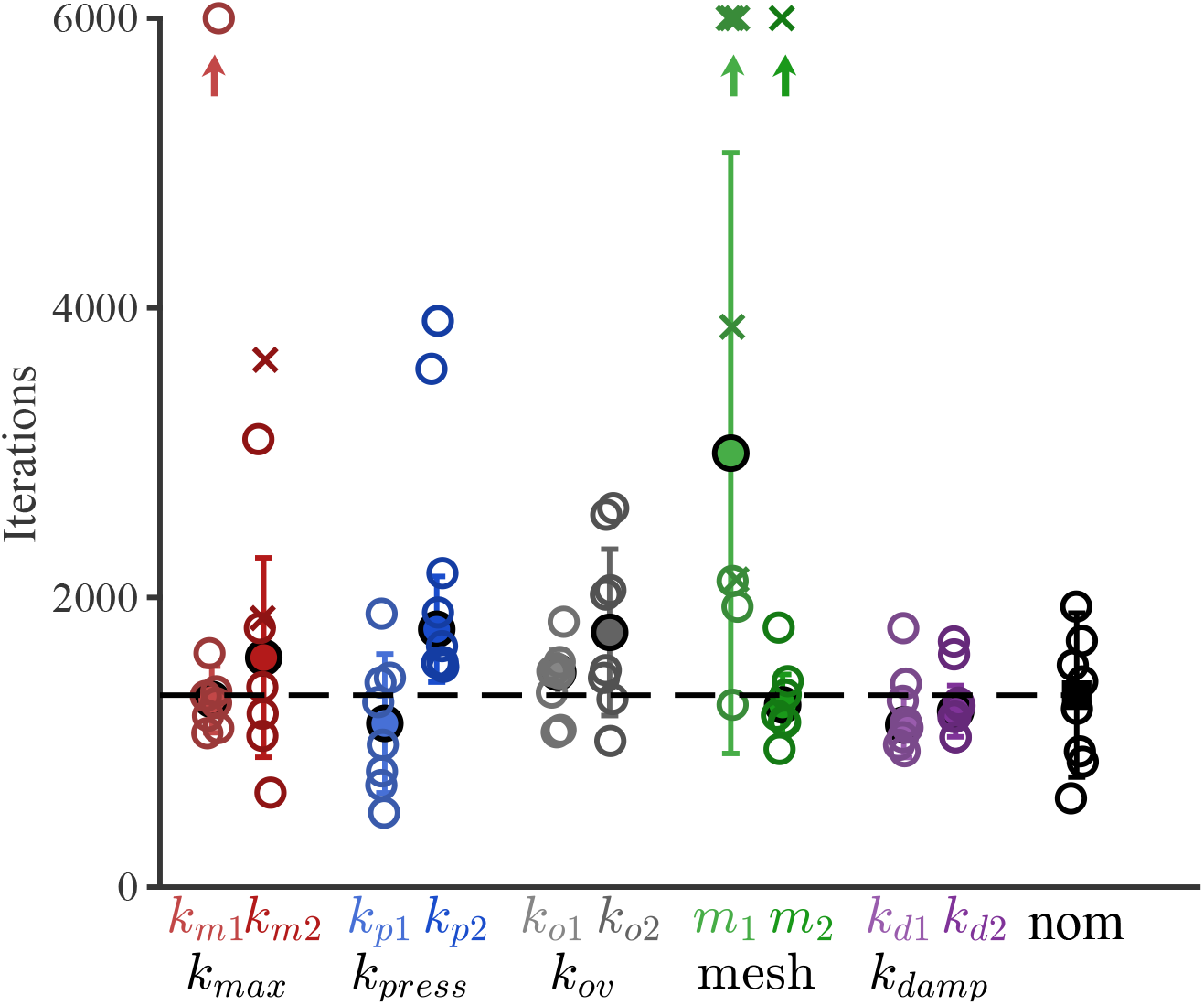
Number of optimization iterations for all parameter configurations. Open circles show individual trials, *×* marks non-converged simulations, filled circles indicate the median (MAD), and arrows denote values beyond the plot limit.

**Fig 3.**
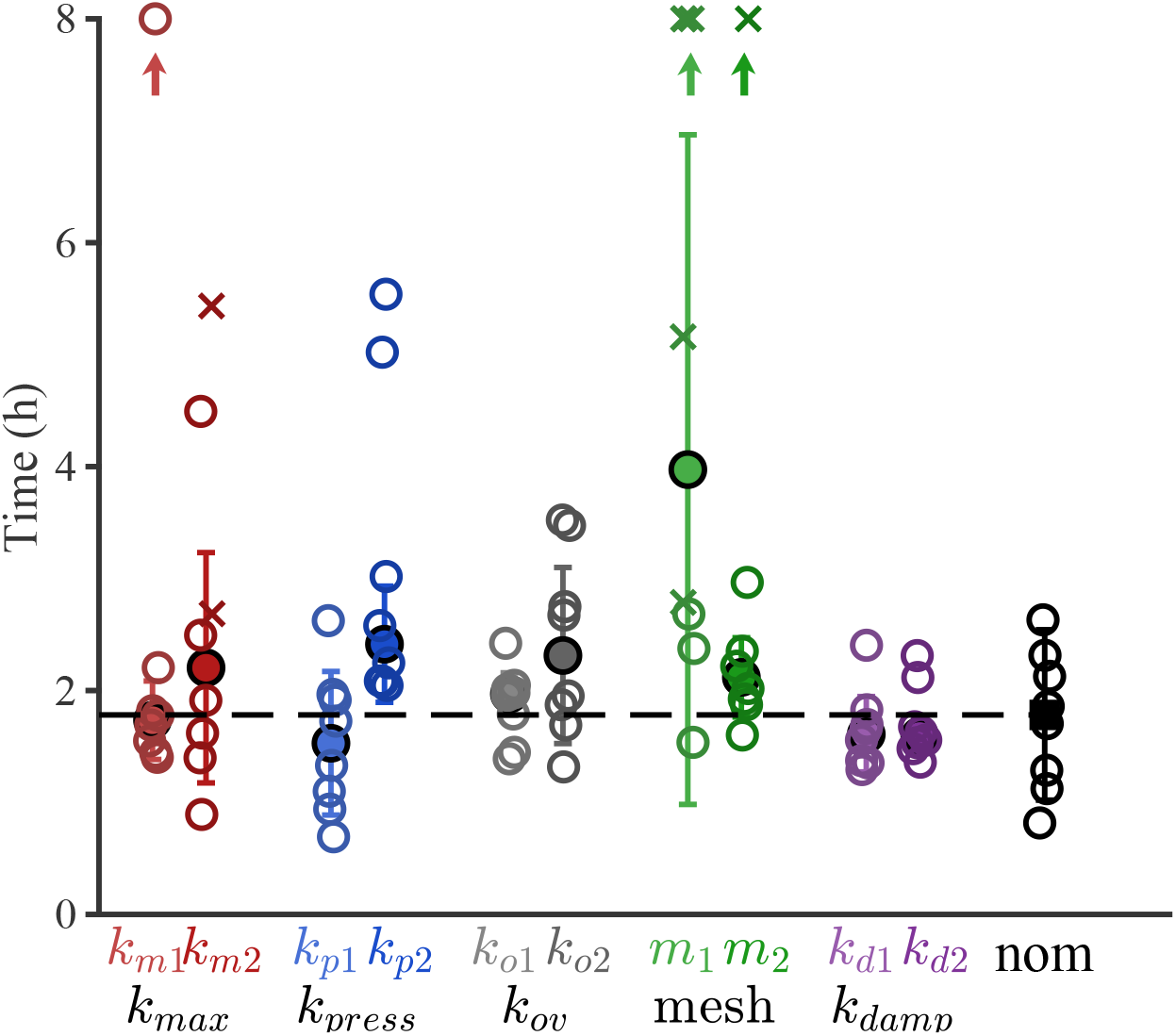
Computation time for all parameter configurations. Each open circle represents one gait trial; crosses (*×*) indicate non-converged simulations. The filled circle marks the median and error bars show the MAD. Upward arrows denote trials that did not converge or exceeded the y-axis limit.

**Table 2.**
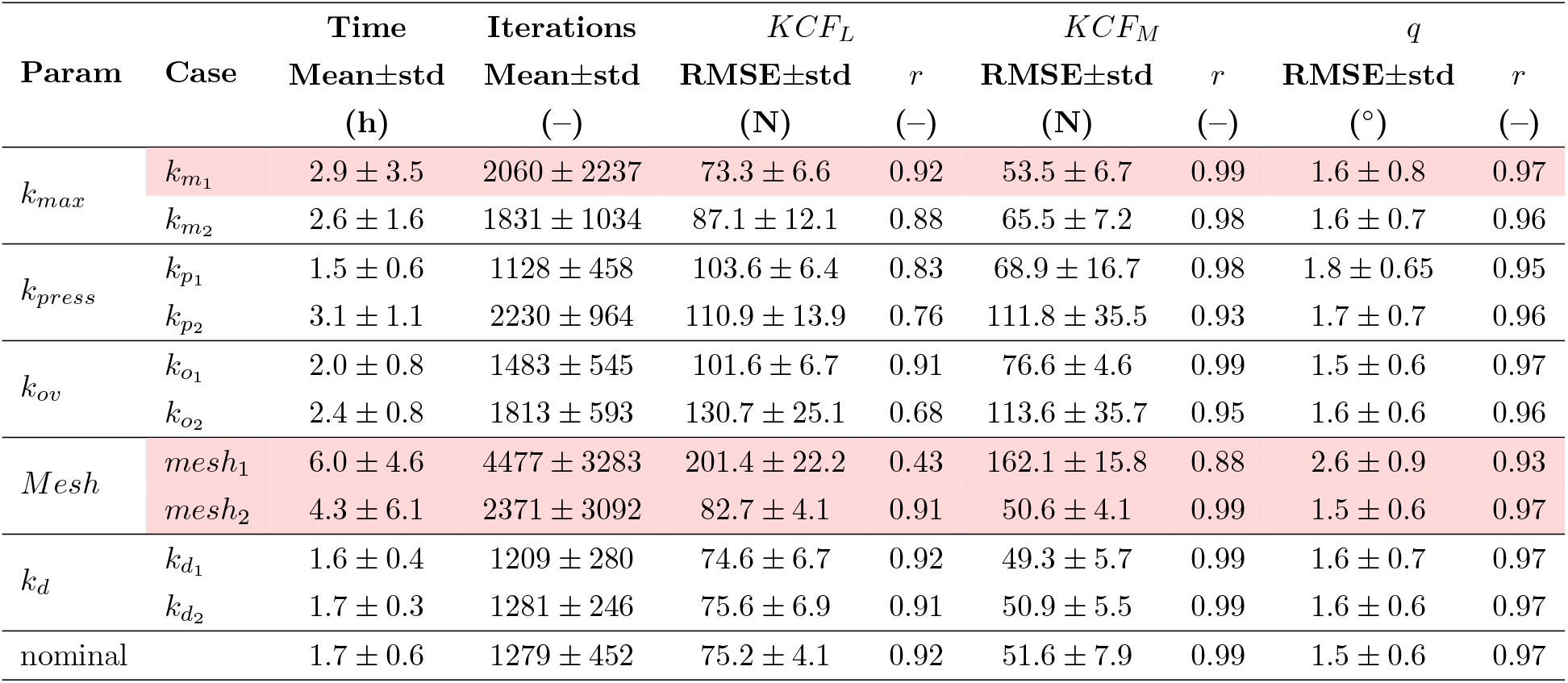
Summary of simulation results for all parameter variations. Values reported as mean *±* std across eight gait trials. Metrics: computation time, iterations, RMSE and *r* for lateral KCF (*KCF_L_*), medial KCF (*KCF_M_*), and joint angles (*q*). Rows highlighted in red contain at least one non-converged trial.

Most parameter combinations applied in the sensitivity analysis converged within 6 hours and fewer than 4000 iterations. Eight simulations did not converge: five were associated with the coarser mesh (*n_tib_* = 49, *n_fem_* = 171), two with the maximum-penetration formulation (non-smooth), and one was associated with the finest femoral mesh configuration (*n_tib_* = 100, *n_fem_* = 342). Most simulations required between 1.5 and 4 hours, while a few parameter combinations reached computation times from 5 to 19 hours (see Fig 3 and Table 2).

The differences in iterations and computational time between the nominal and alternative parameter configurations varied depending on the case, but overall, the stiffer the functions in the contact model, the more iterations were needed to obtain convergence (and more computational time). Mesh resolution had the largest impact on convergence behavior, with both very coarse and highly refined meshes increasing the number of iterations. The use of the coarser mesh led to non-convergence in most simulations (five out of eight). Increasing the smoothing parameter *k_press_* (i.e., reducing model smoothness) also resulted in a higher number of iterations and longer computation times, whereas decreasing this parameter led to slightly faster convergence. Higher values of *k_ov_* and the use of the max function to compute penetration *k_m_*_2_ led to moderately increased convergence cost without improving tracking accuracy. Variations in the knee joint damping parameters had only a minor influence on simulation outcomes, with computation time, number of iterations, knee contact force accuracy, and joint-angle RMSE values remaining similar to the nominal configuration. The finer mesh did not improve significantly the tracking (the RMSE for the medial KCF improved only 1 N in average and the RMSE for the lateral KCF increased 7.5 N in average), and increased computation time and number of iterations (see Fig 4 and Table 2).

**Fig 4.**
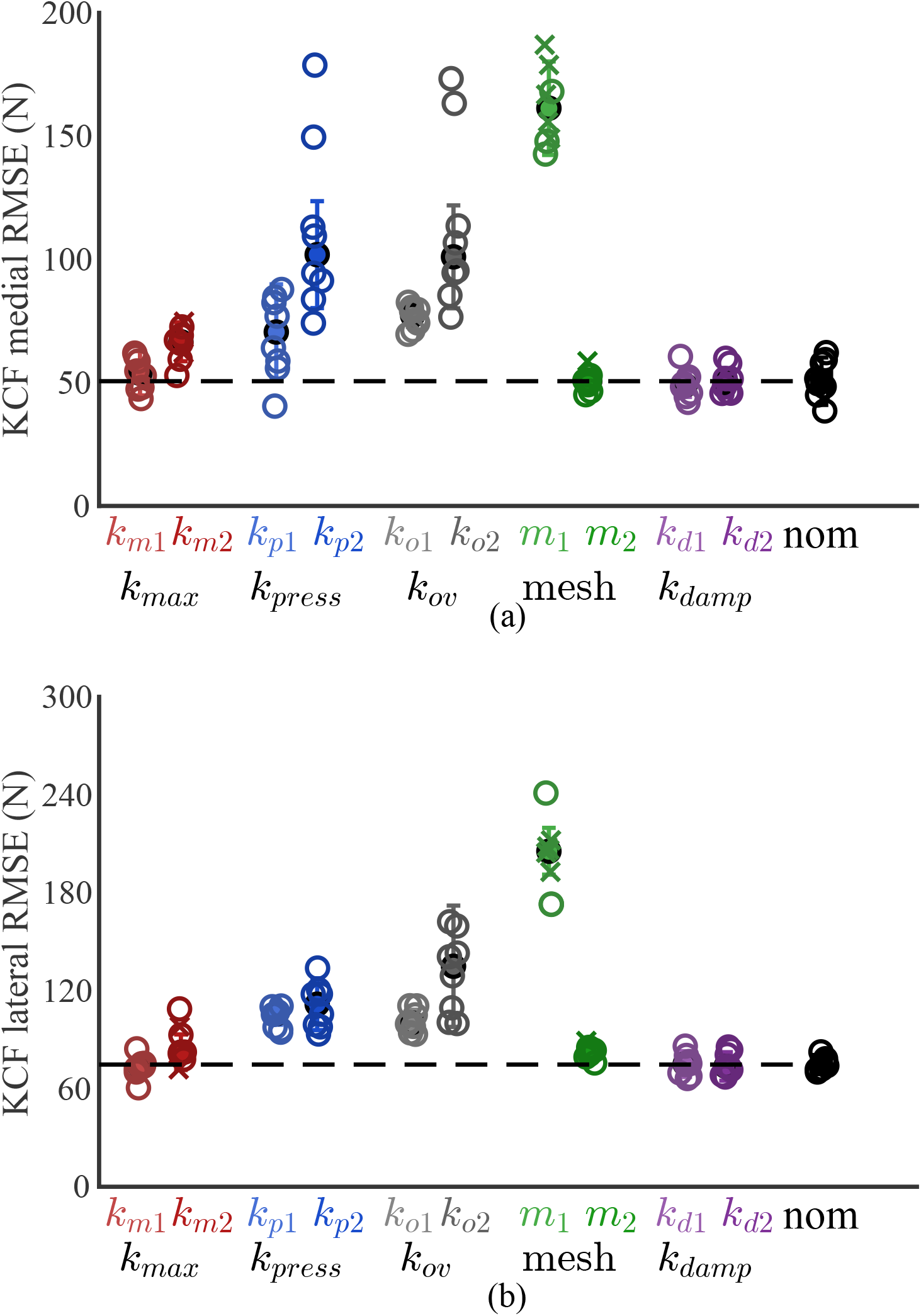
RMSE values of simulated KCFs across all parameter configurations. (A) Medial and (B) lateral KCFs. Each open circle represents one gait trial; filled circles represent the median and error bars the MAD.

Both increasing and decreasing the smoothing parameter *k_press_* overall resulted in higher medial and lateral KCF errors. Using the original *max* function to compute penetrations resulted in slightly lower KCF accuracy, and with *k_max_* = 2 *·* 10^4^ the KCF tracking results were very similar to those obtained with *k_max_* = 10^4^. Larger *k_ov_* values, which produce a stiffer contact transition, led to worse KCF tracking accuracy, particularly for the lateral compartment.

In addition to the resultant knee contact forces, the proposed contact model provides spatial tibiofemoral contact pressure estimates throughout the gait cycle. Contact pressure values were computed for each tibial mesh face at every collocation point of the simulation, providing time-resolved pressure distributions rather than a single resultant contact wrench. The pressure values obtained are summarized as the peak and mean contact pressures for the nominal case for all movements in Table 3. Peak medial tibia pressures ranged from 22 to 28 MPa, while peak lateral tibia pressures ranged from 11 to 17 MPa, depending on the movement task. Mean contact pressures over the entire gait cycle remained below 0.6 MPa in both compartments across all movement conditions. The average RMSE values for joint angles ranged from 1.5*^◦^* to 2.6*^◦^* across all cases (see Fig 5).

**Fig 5.**
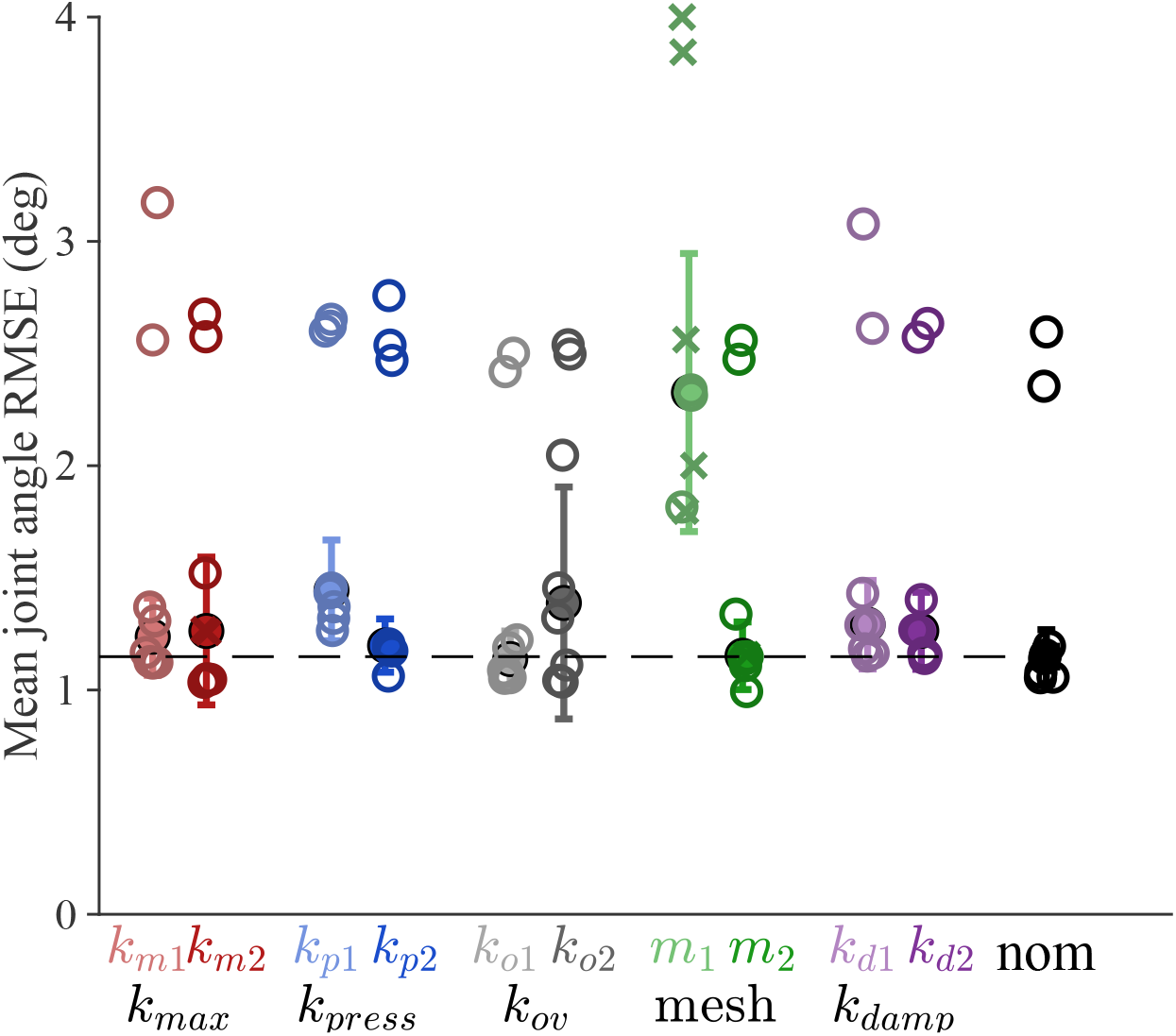
Median joint-angle RMSE values across all parameter configurations. Open circles represent the mean RMSE for each gait trial; the filled circle marks the median across trials; error bars indicate the MAD.

**Table 3.** Pressure peak and average values on the knee artificial joint. Peak and mean tibial contact pressures for the nominal configuration. Peak pressures correspond to the maximum value over all tibial faces and time frames; mean pressures represent the average across the tibial contact surface.

| Movement | Peak Med. (MPa) | Peak Lat. (MPa) | Mean Med. (MPa) | Mean Lat. (MPa) |
| --- | --- | --- | --- | --- |
| bouncy4 | 24.62 | 11.43 | 0.59 | 0.30 |
| bouncy7 | 26.58 | 14.03 | 0.50 | 0.33 |
| mtpgait3 | 22.21 | 13.30 | 0.55 | 0.29 |
| mtpgait9 | 24.78 | 16.83 | 0.51 | 0.34 |
| ngait_og1 | 25.08 | 15.67 | 0.57 | 0.37 |
| ngait_og5 | 26.42 | 14.59 | 0.49 | 0.35 |
| ngait_tm_fast1 | 24.59 | 16.60 | 0.44 | 0.33 |
| ngait_tm_set1 | 28.07 | 10.92 | 0.43 | 0.28 |

The lowest angular tracking errors were obtained for the nominal and *k_o_*_1_ configurations (mean 1.5*^◦^*), followed closely by the refined mesh configuration. The largest joint-angle errors were observed for the coarsest mesh configuration (mean 2.6*^◦^*). Variations in damping and smoothing parameters resulted in slightly increased angular errors relative to the nominal case and did not improve kinematic tracking accuracy.

Overall, the nominal configuration achieved consistently low KCF errors and the lowest joint-angle RMSE values across all gait trials. No alternative parameter configuration provided simultaneous improvement in force accuracy, kinematic tracking, and computational robustness. Consequently, the nominal parameter set was selected as the reference configuration for the predictive simulations.

### Predictive simulations

Predictive simulations were used as a proof of concept to demonstrate the feasibility of incorporating knee contact pressure values into the integral cost function. Increasing the penalization of the maximum contact pressure over the tibial contact surface resulted in a more asymmetrical gait pattern (see S1 Video). The relative durations of the stance and swing phases were similar when no contact pressure penalization was applied (58% stance – 42% swing for the right leg, 54%–46% for the left leg), whereas they diverged as the penalization increased (42%–58% for the right leg, 62%–38% for the left leg), see Fig 6. Higher weighting factors applied to peak knee contact pressure led to lower contact pressure and contact force on both knee compartments, but especially on the lateral side (Fig 7A and 7B, respectively). In turn, muscle contributions to the vertical knee force in the right leg also decreased (Fig 7C). Computational time increased with higher weighting factors: the simulation without penalization required 1.6 h (600 iterations), whereas simulations with *w* = 0.03, *w* = 0.3, and *w* = 3 required 7.7 h (2432 iterations), 17.9 h (6148 iterations), and 17.3 h (6881 iterations), respectively.

**Fig 6.**
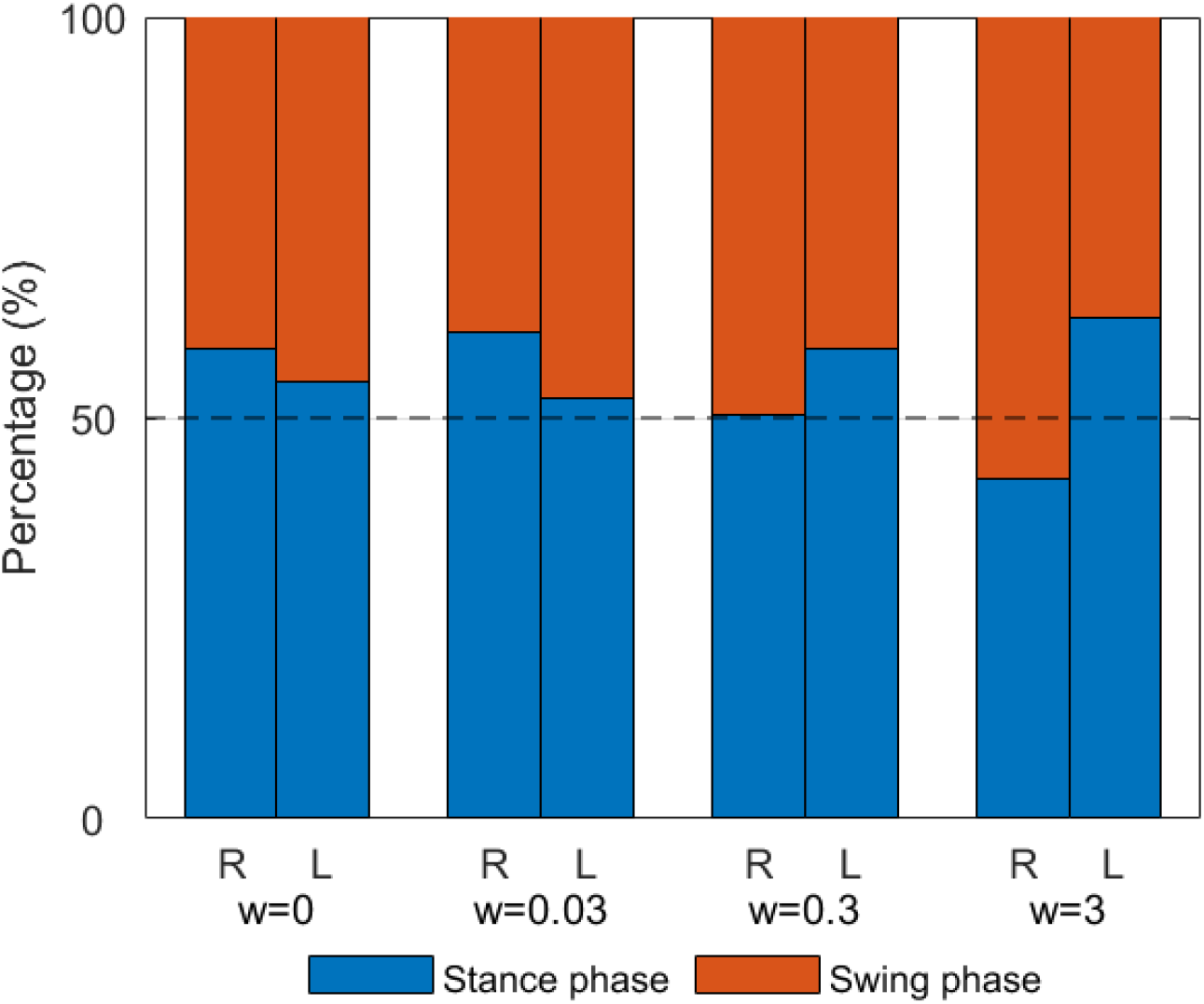
Percentage of the stance and swing phases for the right and left legs, for the four predictive simulations.

**Fig 7.**
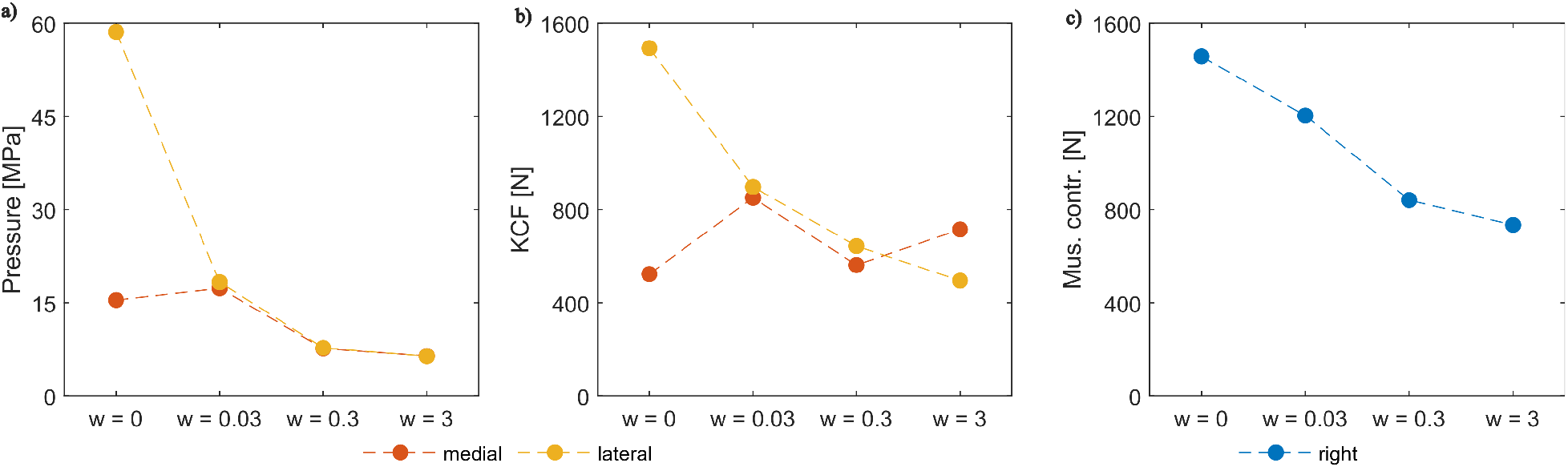
Knee loading after predictive simulations. A) Peak knee normal pressure in the medial and lateral compartments of the right leg; B) peak vertical knee contact forces in the medial and lateral compartments of the right leg; C) peak muscle contributions to vertical knee force in the right leg. Results are shown for four different weighting factors.

## Discussion

This study introduced a smoothed mesh-based knee contact pressure model with a continuously differentiable collision-detection algorithm, enabling its integration into full-body musculoskeletal simulations formulated as OCPs. We present a sensitivity analysis of the key parameters that influence convergence and tracking performance, as well as a proof of concept demonstrating its application in predictive simulations including the values of contact pressures into the cost function.

The results demonstrate that the proposed contact formulation can accurately reproduce experimental lateral and medial knee contact forces (mean RMSE 51.6 N and 75.2 N, respectively) and joint kinematics (mean RMSE 1.5*^◦^*, *r* = 0.97) while maintaining computational efficiency. The nominal parameter set was selected as the reference based on its balanced performance across all evaluated metrics. The nominal case achieved consistently low RMSE values for medial and lateral KCFs and joint angles, and acceptable convergence across all gait trials (as shown in Table 2). While some parameter variations improved individual metrics, none provided simultaneous improvement in force accuracy, kinematic tracking, and computational robustness, supporting the selection of the nominal configuration as an acceptable reference.

Among the parameters examined, the mesh resolution had the strongest influence on both force prediction accuracy and convergence behavior. Overall, mesh refinement only slightly reduced medial KCF error, while increased the average RMSE for the lateral KCF. It also increased computation time and the number of iterations without significant gains in accuracy. In contrast, the coarsest mesh configuration led to decrease knee contact force and kinematic accuracy as well as multiple non-convergent trials. These findings suggest that intermediate mesh resolutions offer the most favorable trade-off between accuracy and computational efficiency.

Smoothing parameters also played an important role in model performance. Increasing the pressure smoothing parameter *k_press_* consistently increased computational cost and KCF error without improving tracking accuracy, indicating that overly stiff contact transitions adversely affect both efficiency and solution quality. Variations in the overlap parameter *k_ov_* primarily affected force accuracy rather than convergence behavior, with larger values increasing RMSE values of lateral KCFs. In contrast, variations in the damping parameters had small influence on simulation outcomes. Across the tested range, damping changes did not meaningfully affect convergence behavior, KCF accuracy, or joint-angle tracking.

The estimated tibiofemoral contact pressures obtained with the proposed formulation for tracking simulations fall within the range reported in previous studies. Peak medial contact pressures reached approximately 28 MPa during dynamic tasks such as bouncy gait, while lateral peak pressures remained consistently lower. These values are comparable to those reported in [11], where the observed peak tibial contact pressures were 22–28 MPa during gait using an elastic foundation-based multibody model. Variation in peak pressure across movement tasks reflects changes in joint loading conditions, whereas mean contact pressures remained low and stable overall.

Unlike Montaut et al. [23], who proposes a randomized smoothing approach to differentiable collision detection for convex shapes, and Beker et al. [24], who propose a smooth analytical formulation of collision detection and contact based on soft signed distance functions, our approach retains a pairwise contact formulation and introduces smooth approximations for both penetration computation and false-positive contact detection. Similar elastic foundation-based approaches have been used in musculoskeletal simulations estimating muscle activations and kinematics simultaneously alongside the tibiofemoral cartilage contact pressures during dynamic tasks; however, these methods were developed for inverse approaches (i.e., when ground reaction forces and most kinematics variables are known) [13, 14, 25, 26]. In line with these methods, the present formulation enables pressure estimation within full-body tracking. The main novelty is that it is continuously differentiable and can be integrated into full-body predictive simulations, allowing contact pressures to be incorporated into the integral cost function.

As a proof of concept, the model was incorporated into predictive simulations, demonstrating that pressure-based cost terms can drive gait adaptations that reduce tibiofemoral loading, particularly in the lateral compartment. The results showed that higher penalization of the peak joint pressure decreased the peak contact pressure value and subsequently reduced muscle contributions to the knee inferior-superior force, as expected [27]. The present validation was performed on a single subject with a knee prosthesis, and extension to native cartilage geometries remains an important next step.

The main limitations of this study relate to the modeling of passive forces. In this study, passive moments were introduced at all joints, as well as passive moments and forces at the knee degrees of freedom. These quantities increase exponentially near the limits of joint ranges of motion (as in [17]). These are intended to represent the passive contribution of ligaments; however, a more detailed representation of the knee ligaments would likely improve the accuracy of both contact and muscle force estimates [28]. Similarly, the number of faces used here is relatively low compared to the full prosthesis geometry. However, the sensitivity analysis demonstrated that the nominal mesh resolution performed comparably to the finer configuration in terms of KCF and kinematic tracking. A finer mesh resolution would become more relevant when combined with a more detailed representation of passive structures, such as ligaments and cartilage, which would improve the accuracy of both the contact pressure distributions and passive force estimates.

Caution is also required when interpreting predictive simulations that do not use experimental KCF data as input. Kinematic results differ only slightly between tracking simulations, which use experimental data as input, and predictive simulations without pressure penalization, which do not rely on such data. In contract, KCF estimates differ substantially between the two approaches. This observation is consistent with previous studies showing that predicted knee contact and muscle forces are sensitive to modeling assumptions and parameter choices that are often difficult to calibrate experimentally [29]. Moreover, discrepancies between predictive simulation results and measurements may arise from both biomechanics modeling aspects (e.g., the formulation of physiological relationships and the identification of model parameter values [22] and the definition of cost function terms [17]. A potential approach to mitigate these discrepancies is to solve inverse optimal control problems. Future work will also focus on extending the model to natural knee geometries with cartilage representation, multi-subject validation, and clinical applications such as osteoarthritis assessment and evaluating the effects of assistive devices on pain and joint loading.

This model enables the study of contact pressures within reasonable computation times on standard computers and in a range of application scenarios, such as evaluating the effects of assistive devices on human performance and pain, or investigating determinant factors of osteoarthritis in human joints.

## Conclusion

This paper presented a smoothed mesh-based knee contact model that computes continuously differentiable tibiofemoral contact pressures within full-body musculoskeletal simulations formulated as OCPs. By introducing smooth approximations for maximum penetration detection, contact pair validation, and pressure computation, the proposed model overcomes the non-smoothness limitations of applying mesh-based collision-detection algorithms, enabling its direct integration with gradient-based solvers and automatic differentiation tools. This feature enables new studies, including ‘what if’ scenarios in which knee contact pressure is treated as an objective to be optimized by the central nervous system.

## Supporting information

**S1 Video. Predictive simulation gait adaptation.** Animation showing the asymmetric gait pattern generated by predictive simulations with increasing knee contact pressure penalization weights (*w* = 0, *w* = 0.03, *w* = 0.3, *w* = 3).

## A Smoothing functions

### A.1 Smoothing of the maximum penetration value

The Mellowmax function (Eq 1) acts as a maximum function when *k*_max_ *→ ∞* and as a mean value when *k*_max_ *→* 0. As the smoothing parameter increases, the result becomes closer to the true maximum value, but the function is less smoothly differentiable. Conversely, as the parameter decreases, the function becomes smoother but deviates more from the true maximum value.

Fig 8A illustrates the penetration values between one tibial face and all potential femoral faces when rotating the knee in flexion-extension from 0*^◦^* to 90*^◦^*, while keeping all other DoFs fixed. The maximum values of these curves are shown in the black curve of Fig 8B. Note that the first derivative 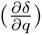 would not be continuous. Red and yellow curves show the maximum values when using the Mellowmax function with *k_max_* of 10^4^ and 2 *·* 10^4^, respectively. The shadowed area shows the region plotted in Fig 8C as a zoom.

**Fig 8.**
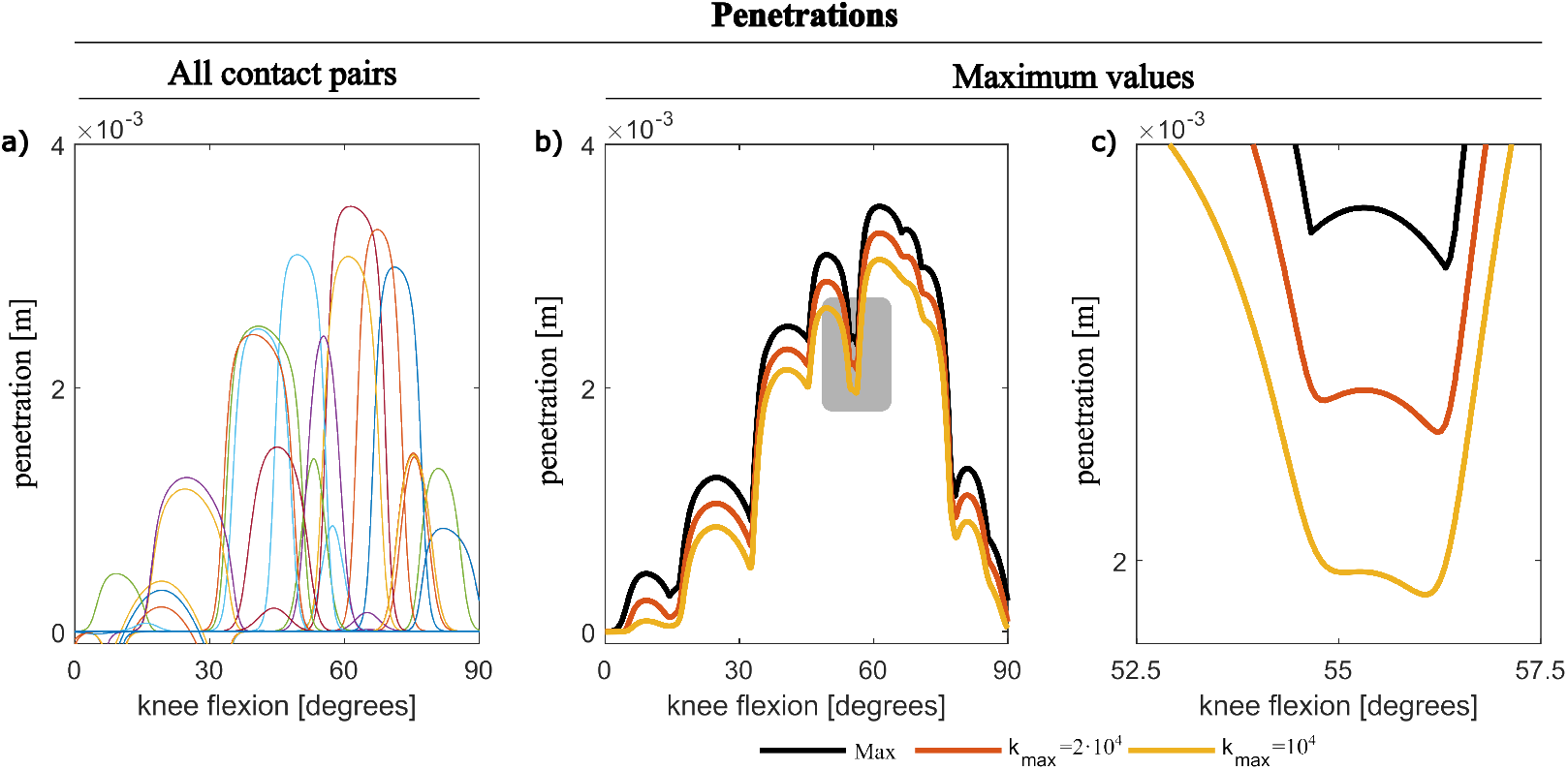
Example of smoothing the maximum penetration value. A) Penetration of one tibial face *δ_i_*against all candidate femoral faces. B) Maximum of the curves in (A), computed using the exact maximum function (black) and Mellowmax function approximation with *k_max_* = 10^4^ (red) and 2 *·* 10^4^ (yellow). C) Zoomed-in view of the shaded region in (B), illustrating the smoothing effect.

### A.2 Smoothing of the contact pressure transition

The smooth pressure transition defined in Eq 4) is controlled by the parameter *k_press_*. The penetration *δ* is negative when the two surfaces are separated and positive when they are in contact.

Panel (A) of Fig 9 shows the pressure-penetration curve for three different values of *k_press_*. The transition was shifted to *δ*_0_ = 0.3*mm*, rather than being ceneterd at zero, to reduce the magnitude of non-physical negative pressures for negative penetration values. With a small value of *k_press_*, such as 10^3^, the transition is very wide and the pressure values remain non-negligibly negative over a substantial range. Although this behavior is smooth, it is not desirable, because negative pressures are not physically meaningful. With *k_press_* = 10^4^, the transition becomes narrower and more accurate with respect to reference curves. The pressure still changes smoothly around *δ*_0_, and the negative pressure values are negligible. With a high value such as 10^5^, the transition is extremely sharp. Panel (B) zooms in on the region from *−*0.5 to 0.5 mm: the small value (10^3^) clearly shows a wide and soft transition with negative pressures; the intermediate value (10^4^) produces a smoother but more accurate transition; the highest value (10^5^) limits the change to a very narrow region around *δ*_0_. Because the transition uses a tanh-based function, the curve remains smooth for any finite *k_press_*, avoiding sudden jumps in pressure during simulations.

**Fig 9.**
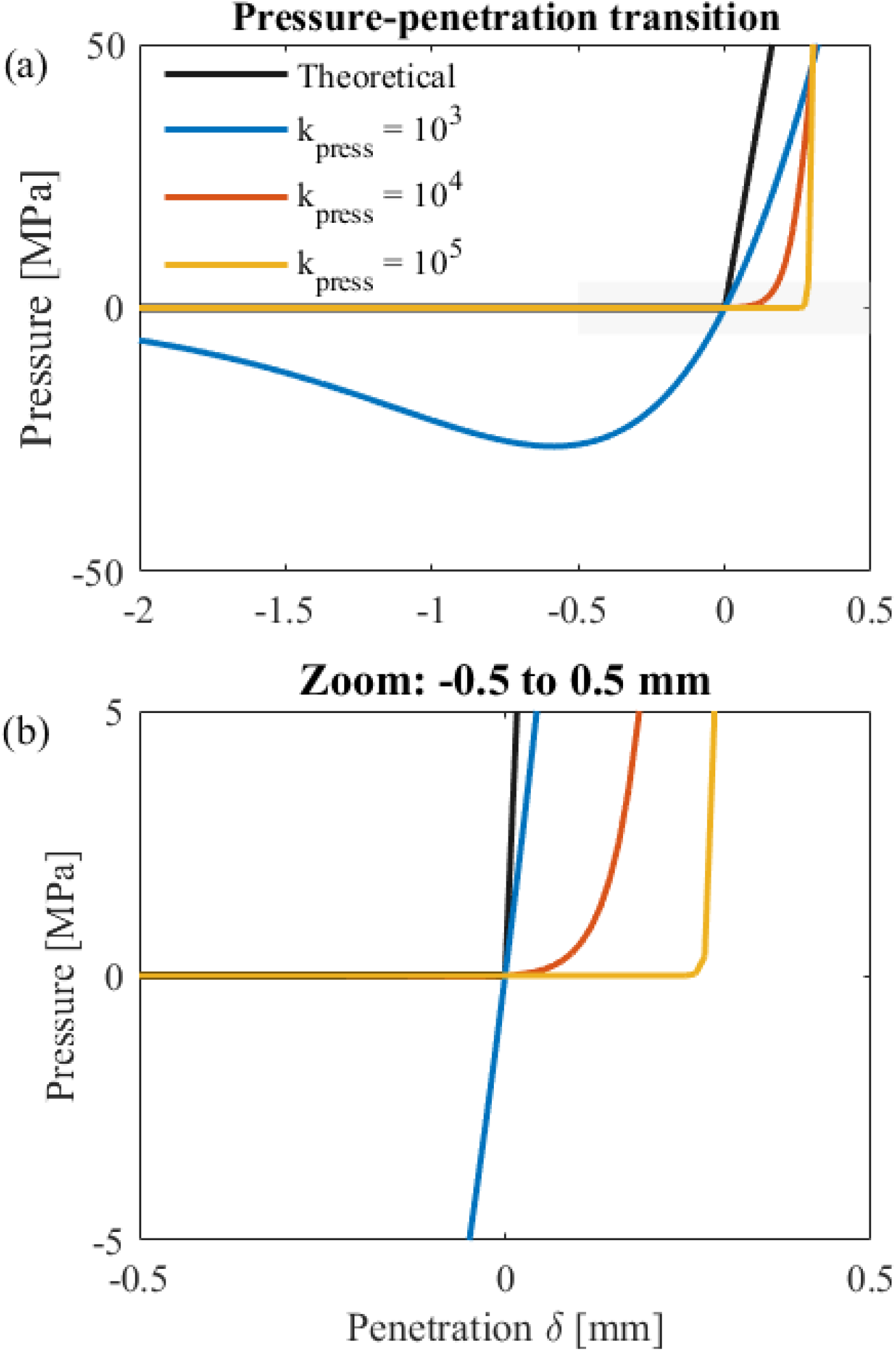
Effect of the smoothing parameter. *k_press_* **on the pressure-penetration relationship.** (A) Pressure as a function of penetration *δ* for three smoothing values (10^3^, 10^4^, and 10^5^). (B) Zoomed view of the region *−*0.5 to 0.5 mm, highlighting how increasing *k_press_* narrows and sharpens the transition around *δ*_0_.

### A.3 Contact weighting factor

To avoid false contact pressures, we check whether the orthogonal projection *P_i_* of the tibial-face centroid **c***_i_*onto the plane of femoral face *j* (along the tibial normal 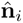) lies within a footprint circle defined for that femoral triangle. For each femoral triangle *j*, we define a center **o***_j_* and threshold radius *d*_thres*,j*_ as follows: if the triangle is acute, **o***_j_* is the circumcenter and *d*_thres*,j*_ is the circumradius; if the triangle is obtuse, **o***_j_* is the midpoint of the longest edge and *d*_thres*,j*_ is half the length of that edge.

If *P_i_* lies within the circle centered at *o_j_* with radius *d*_thres_ (i.e., *d*_mult_ = ||*P_i_* − o_j_|| ≤ *d*_thres_), we retain the pair as a potential contact; otherwise, we discard it. In practice, we replace this hard threshold with a smooth multiplier *mult ∈* [0, 1] based on the distance *d*_mult_ = ||*P_i_ −* **o***_j_*||:

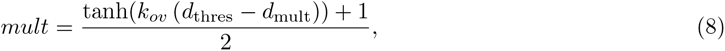

where *k_ov_* is a smoothing parameter. The behavior of the smooth multiplier can be understood by introducing *s* = *d*_thres_ *− d*_mult_, which is positive when the projected point is within the threshold. As *s* increases, the multiplier moves smoothly from 0 to 1. Fig 10A shows this transition for different values of *k_ov_*. With *k_ov_*= 100, the transition from 0 to 1 is very smooth. With *k_ov_* = 1000, the transition becomes sharper, and with *k_ov_* = 10 000, it occurs over less than 0.5 mm. Fig 10B provides a zoomed view of the region around *s* = 0, highlighting how the smoothing parameter controls the width of the transition band. Because the function in Eq (8) uses a hyperbolic tangent, the curve and its derivative remain smooth for any finite *k_ov_*, preventing abrupt changesduring optimization.

**Fig 10.**
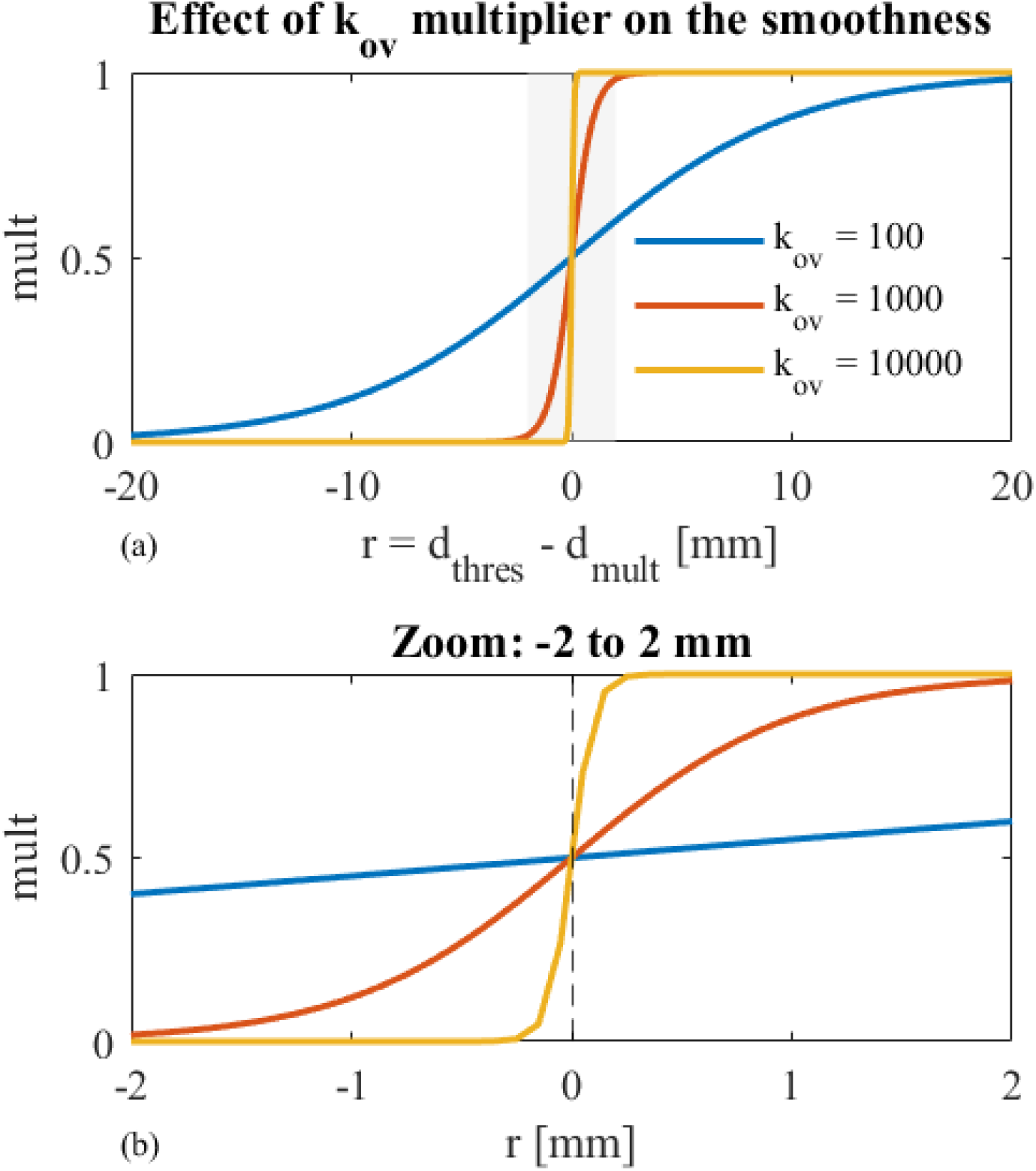
Effect of the smoothing parameter *k_ov_* on the multiplier. (A) Multiplier as a function of *r* = *d*_thres_ *− d*_mult_ over a wide range of distances, for *k_ov_* = 100, 1000, and 10,000. (B) Zoomed view around *r* = 0 (from *−*2 to 2 mm), showing how small *k_ov_* values produce a more gradual transition, whereas larger values result in a steeper and more localized transition from 0 to 1.

